# A multiscale physicochemical characterization of melanin-based feather coloration in the rapid Capuchino Seedeater radiation

**DOI:** 10.1101/2025.10.29.685391

**Authors:** Stephan M. Sutter, Olivia G. Rogers, Daniel L. Bilezikian, Casey B. Dillman, Teresa J. Porri, Luís Fábio Silveira, Sheela P. Turbek, Lara A. Estroff, Leonardo Campagna

## Abstract

Melanin-based plumage coloration in birds is shaped by pigment composition as well as melanosome morphology and distribution, however, the ways in which these factors together modulate observable color remain poorly understood. We investigate this relationship in the Capuchino Seedeaters (genus *Sporophila*), whose recent, rapid radiation driven by sexual selection resulted in 12 species with diverse coloration patterns. Using scanning electron microscopy (SEM) and micro-computed tomography (µCT), combined with a novel application of Fontana-Masson stain to image melanosomes at high resolution, we characterize melanosome distribution and morphology in several variably colored plumage patches across Capuchino species. Melanosome morphologies followed patch-specific patterns that did not directly correlate with coloration: crown feather melanosomes were larger, more elongated, and had greater percent eumelanin content than those in belly, throat, or dorsum/rump patches. We also observed that dorsal patches had more total melanin than ventral ones, with pigment and coloration patterns suggesting possible signaling and photoprotective roles. More generally, we show how the patch-specific coloration of male Capuchinos is accompanied by differences in melanosome morphology and melanin composition and abundance. Our work highlights the challenges that remain in understanding how the nanoscale mechanisms of melanin-based pigmentation translate into macroscale plumage coloration.

## 1. Introduction

The wide diversity of avian plumage coloration arises from two principal mechanisms: the physical interaction, *e*.*g*., diffraction or refraction, of light with the internal barb and barbule keratin microstructure (structural color), and the unique absorption profile of one or more feather pigments (pigmentary color) [1]. The most ubiquitous pigments include melanins, which are endogenously synthesized (and thus genetically controlled) and produce muted, black or dark brown to reddish coloration [2–6]; and diet-derived carotenoids, which are responsible for bright red, orange, and yellow plumage [4,5,7,8]. Unlike structural color, in which precise nanoscale structures cause optical interference that can often be modeled with high fidelity [9–15], pigmentary color is controlled by a hierarchy of factors from the highly conjugated structures of pigment molecules [6,16–18], to their encapsulation within melanosomes (in the case of melanin; Fig. 1) [15,19], and ultimately their spatial distribution within the feather [19–22]. Analysis of pigment content often cannot fully account for observed color [23–27], and efforts to understand the link between melanin deposition and the macroscale feather phenotype have yet to conclusively demonstrate the ways and extent to which combinations of pigment type(s), melanosome morphology, and pigment distribution contribute to plumage coloration. Here we use a combined, multiscale analysis of pigment chemistry and melanosome morphology to evaluate the role of melanosome morphology on plumage coloration in the Capuchino Seedeaters [28,29], a cohort of 12 tanager species (genus *Sporophila*) that resulted from recent speciation driven by sexual selection [30–32].

**Figure 1.**
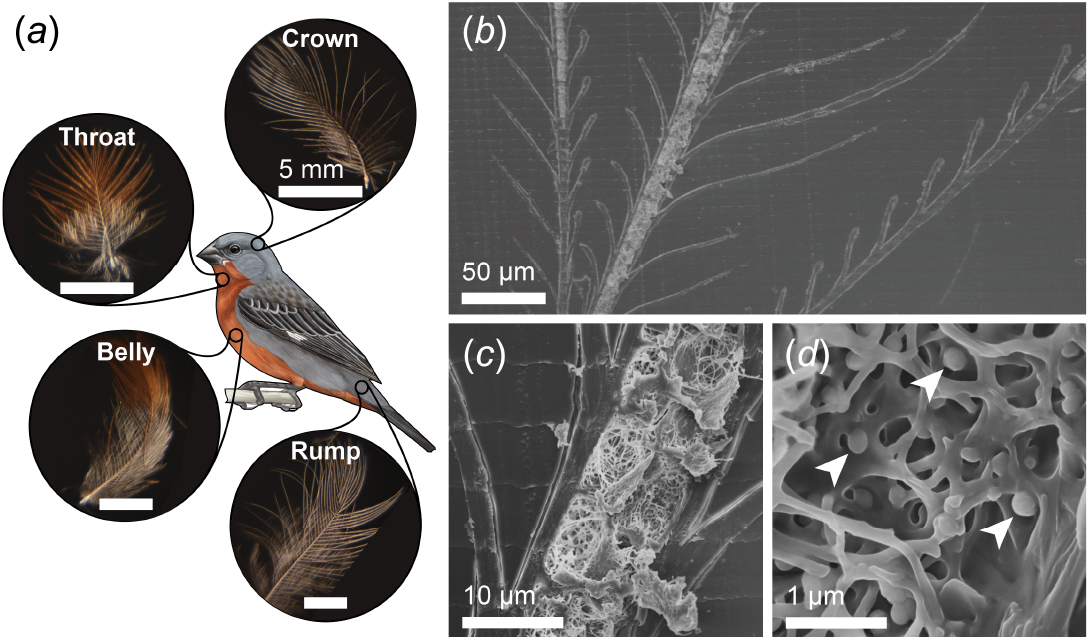
Overview and examples of a *Sporophila* Seedeater’s feather patches and feather microstructure. (*a*) Representative feathers (optical images, *left*) of *S. castaneiventris* feathers from different body patches: crown, throat, belly, and rump. In this species, the crown and rump patches are both gray, and the throat and belly patches are both cinnamon. (*b*) Secondary electron SEM image of an embedded and sectioned *S. iberaensis* feather showing three exposed barbs, each with many branching barbules. (*c, d*) The interior of a sectioned barb, showing the keratin matrix and several round melanosomes (white arrows).

Among the melanins, a structurally diverse family represented across every kingdom of life, eumelanin (EM) and pheomelanin (PM) are the two canonical integumentary pigments that are broadly conserved in birds, mammals, and certain fish and reptiles [1]. Both EM and PM have highly conjugated structures derived from a shared tyrosine precursor [16,18,33], but their respective synthesis pathways and subsequent organization within melanosomes give rise to differing nano- to macroscale properties. Melanin biosynthesis occurs during early feather development within melanosomes, small spherical to ellipsoidal organelles that remain trapped within the rigid keratin matrix of the mature feather [34]. EM synthesis occurs *via* polymerization of 5,6-dihydroxyindole (DHI) and DHI-2-carboxylic acid (DHICA) subunits on scaffolds of the fibrillar protein PMEL17 [35], which provide eumelanosomes the internal rigidity to accommodate a range of low- to high-aspect ratio (*i*.*e*., ellipsoidal) morphologies. The resulting pigment has a broad absorption band spanning most of the visible spectrum and into the ultraviolet (UV) [2], giving EM its characteristic dark brown to black appearance. PM is composed of varying proportions of benzothiazine (BT-PM) and benzothiazole (BZ-PM) moieties [16], which have lower absorption at visible wavelengths (particularly in the range of 550–650 nm) and produce red to orange or yellow hues [36,37]. The downregulation of PMEL17 during PM synthesis results in a spherical melanosome morphology [38].

The macroscale coloration that melanosomes confer to a feather can be influenced by both their pigment content and their morphology: the encapsulation of pigment within melanosomes adds a level of complexity by allowing different melanosome configurations to produce a range of drab to iridescent feathers [15,19,39,40]. Recent discoveries of preserved melanosomes in the fossilized integumentary structures of feathered dinosaurs [40–46], prehistoric birds [47–50], and other extinct species [51–55] have motivated efforts to correlate melanosome morphology and distribution [56] with macroscopic feather appearance, using extant bird species as a reference to reconstruct the coloration of fossilized tissues. A major confounding factor is the absence of straightforward, generalizable correlations between melanosome morphology and feather color, making it difficult to impute pigment content and feather color directly from melanosome morphology [40,43,45]. Data analysis methods like quadratic discriminant analysis can nevertheless enable accurate prediction of some feather colors on the basis of morphological parameters [40,43,51,57,58] using multi-species datasets such as the one in Li *et al*. (2010) and Li *et al*. (2012), which have strong predictive power for black, brown, and gray feathers [40,43]. Nevertheless, species- and patch-specific heterogeneity in pigment content, melanosome morphology, feather microstructure, and macroscopic color poses a challenge to generalizing such findings, and highlights the complex relationship between melanosomes and plumage coloration.

The physicochemical interactions of melanin and melanosomes with UV light contribute to important roles in structural color, photoprotection, and signaling. Most birds have at least some feathers with strong UV reflectance arising from either pigmentary or structural coloration [59–62], the latter occurring paradoxically in iridescent colors derived from the periodic ordering of melanosomes (the contents of which otherwise strongly *absorb* UV) [15,19,63–66]. Both EM and PM release radical species through UV-induced photodegradation [8,67], but EM does so to a much lesser degree and is associated with a greater photoprotective effect [37,68–70]. The partitioning of strongly UV-reflective feathers to small, distinct anatomical regions suggests alternative roles in signaling through contrast between differentially reflective patches [1,60,71,72]. Indeed, the UV reflectance of plumage is no less important than its visible color, as a UV-sensitive opsin variant gives birds tetrachromatic vision and the ability to differentiate plumage colors beyond the visible spectrum into the UV-A range [73–75].

Assessing the different contributions of the keratin matrix, melanosomes, and pigment content to feather color requires the use of multiscale, complementary characterization techniques. Structural keratin motifs occur on length scales in the range of tens of microns, and can therefore be directly visualized by scanning electron microscopy (SEM) [25,76–78] or micro-computed X-ray tomography (μCT) [76–81]. μCT permits nondestructive three-dimensional imaging, but its limited resolution makes it more suited for studying mesoscale matrix structures and barb and barbule geometries; scanning and transmission electron microscopy (SEM and TEM, respectively) can resolve melanosomes and other structures on the scale of hundreds of nanometers to microns, but require complicated sample preparation [15,19,21,65,82,83]. Specific (*e*.*g*., immunogold) or nonspecific (*e*.*g*., lead citrate) labels are often used in TEM to improve the contrast of biological structures [84] including melanosomes [80,81], however, such stains are not typically employed in SEM. Individual melanin pigment molecules are too small to visualize, but can nevertheless be characterized using UV-visible spectroscopy [59,63] as well as high-performance liquid chromatography (HPLC) and mass spectroscopy [85,86]. Integrating structural and pigment datasets remains challenging: while electron microscopy permits morphological analysis across a range of length scales down to the single-melanosome level, albeit with low throughput, the detection limits of biochemical and optical techniques typically necessitate bulk measurements over a macroscopic portion of a feather. As a result, direct links between pigment composition, melanosome morphology, and feather color remain poorly understood.

Here we examine the relationships between both the melanin content and morphological characteristics of barb melanosomes and observable plumage coloration, using the Capuchino Seedeaters as a model system. The Capuchinos are 12 species of the genus *Sporophila*, a group of Neotropical tanagers that underwent rapid speciation during the Pleistocene with most species diverging within the last million years [87] in the subtropical and temperate grasslands of modern-day Brazil, Paraguay, Uruguay, Argentina, and Bolivia [28,29]. Genomic comparisons suggest that the Capuchino radiation was driven by sexual selection through female mating preference primarily based on male plumage color and song [30–32]. The Capuchinos consequently show strong sexual dimorphism: males can be distinguished by their vocalization as well as unique patterns of black, gray, cinnamon, rufous, cream, and white-colored plumage patches that emerge seasonally during the austral summer mating period, while females of all species have similar drab, olive-brown coloration [75,88]. Notably, interspecific differences in genes associated with melanin biosynthesis and melanosome trafficking that give rise to this phenotypic variation constitute most of the genomic divergence found between Capuchino species [23,31]. The Capuchinos therefore provide a unique and tractable opportunity to study the physicochemical mechanisms behind melanin-based feather coloration, as they allow comparisons between several plumage patches and colors, within and between closely related species differing in few other traits.

Building on recent work linking differences in EM and PM content to feather color variation in Capuchinos [23], we combine melanin-specific staining with scanning electron microscopy (SEM; Fig. 1) and micro-computed tomography (μCT) to characterize the melanosome morphologies seen across different feather colors, patches, and species. We hypothesize that melanosome morphology may vary to accommodate differences in pigment content, and contribute to the finer-scale differences between similar hues (*e*.*g*., black vs. gray, or cinnamon vs. rufous) not explained by pigment quantification. Our findings identify differences in the production of the six patch-specific feather colorations (cinnamon, rufous, black, gray, cream, and white), and show that variation in pigment type and concentration are accompanied by changes in melanosome size and shape. Additionally, we show that similar plumage colors can arise from different melanosome morphologies, complicating efforts to predict coloration from melanosome shape alone and underscoring the importance of multiscale characterization in the investigation of color production mechanisms.

## 2. Materials and Methods

### 2a. Feather collection

Our sample set included feathers from the crown, dorsum/rump, throat, and belly patches (Fig. 1) of the 12 Capuchino species: *Sporophila bouvreuil, S. cinnamomea, S. hypochroma, S. hypoxantha, S. iberaensis, S. melanogaster, S. nigrorufa, S. palustris, S. pileata*, and *S. ruficollis*, comprising the core radiation or southern Capuchinos sensu Lijtmaer et al. (2004) [29]; as well as the closely related northern Capuchinos *S. castaneiventris* and *S. minuta*. All feathers were collected at the peak of the Capuchino mating season (austral summer), as males undergo seasonal cycles between brightly colored mating plumage and dull eclipse coloration. We obtained feathers from archived collections at the Centro de Ecología Aplicada del Litoral (CECOAL; Corrientes, Argentina) and the Museu de Zoologia da Universidade de São Paulo (São Paulo, Brazil). A full list of the feathers we sampled can be found in Table S1.

### 2b. Melanin content quantification

The quantification of melanin content was previously reported by Estalles *et al*. (2022) and included, for each patch of 10 southern Capuchino species, the mean EM and PM content in micrograms of pigment per milligram of feather mass [23]. These were calculated in Estalles *et al*. (2022) by quantifying the hydrolysis products of EM (pyrrole-2,3,5-tricarboxylic acid; PTCA) as well as BT-PM (4-amino-3-hydroxyphenylalanine; 4-AHP), and BZ-PM (thiazole-2,4,5-tricarboxylic acid; TTCA) moieties by HPLC, and applying correction factors (38, 34, and 9, respectively) to infer the corresponding pigment content (Table S2, [23]).

### 2c. Feather washing

To remove debris and disentangle barbs, we first washed and dried feathers using a protocol adapted from Igic et al. (2018) [25]. We agitated the feathers in a 5% (w/v) dilution of standard dish soap (Dawn) in reverse-osmosis (RO) water for 15 seconds, followed by a 30-second wash with RO water. We subsequently washed the feathers twice for five minutes in 100% ethanol (Koptec Pure Ethanol, 200 proof), followed by a final five-minute wash with RO water, after which we gently dried the feathers with compressed nitrogen.

### 2d. Whole-feather Fontana-Masson staining

Fontana-Masson (FM) staining is a common histological technique that takes advantage of the ability of melanins to selectively reduce aqueous silver ions (Ag^+^), forming insoluble elemental silver (Ag^0^) deposits where melanin is present. The dark brown color of melanin-bound silver, further toned (darkened) with a thin surface layer of gold, conventionally allows the pigment to be identified in mounted tissue sections through optical microscopy, but the ability to selectively label melanin with a heavier element gives FM a new use as a positive stain for back-scattered electron scanning electron microscopy (BSE-SEM). Here, FM staining of whole feathers was performed by the Cornell University College of Veterinary Medicine Animal Health Diagnostic Center using a standard FM protocol for argentaffin staining of tissue sections, modified only by exclusion of the unnecessary de-paraffinization step (see protocol in supplemental information).

### 2e. Resin-embedding feathers for SEM imaging

We resin-embedded feathers in a two-step process [89]. In the first step, we removed the proximal (plumulaceous) section of each feather and laid the remaining (pennaceous) section flat in a shallow, 20×20×1 mm silicone monolayer mold (Electron Microscopy Sciences, Hatfield, PA) that we then flooded with EMbed 812 resin (Electron Microscopy Sciences), using a toothpick to remove any bubbles. We then cured this first embed at 60°C for 24 hours. In the second step, we trimmed the square resin block containing the feather, to fit standing vertically at one end of a rectangular silicone embedding mold (14ξ5ξ4 mm; Electron Microscopy Sciences). We then filled the rest of the mold with resin and cured it at 60°C for 24 hours, yielding a rectangular resin block with the feather embedded at one end (parallel to the smallest face of the block) to permit longitudinal sectioning.

### 2f. Embedded feather sectioning and preparation for SEM imaging

We obtained exposed longitudinal sections of the interior keratin matrix of each feather using an Ultracut UCT ultramicrotome (Leica Microsystems, Wetzlar, Germany) with a freshly cleaved glass knife. We mounted each resin block in the microtome chuck such that the cutting direction ran parallel to the rachis of the embedded feather. After roughly facing the resin block, we made cuts in 2–5 μm increments until a suitable longitudinal section through one or more barbs was exposed on the block face.

### 2g. Scanning electron microscopy

Secondary-electron SEM (SE-SEM) is often used to visualize the nanoscale morphology of biological structures, while backscattered-electron SEM (BSE-SEM) provides contrast based on atomic number (Z), with heavier elements scattering more strongly and appearing brighter. To visualize melanin we leveraged Fontana-Masson (FM) staining, in which heavier melanin-bound silver (Z = 47) and gold (Z = 79) produce strong BSE contrast against the lighter elements (e.g. carbon, nitrogen, oxygen) of the surrounding keratin. Accordingly, we used SE-SEM to image melanosome morphology and BSE-SEM in combination with FM staining to selectively visualize melanin deposits against the unstained keratin matrix. Samples were sputter-coated with either Au-Pd at 20 mA for 45 s, or carbon at 40 A for 30 s, using a Desk V sputter coater (Denton Vacuum, Moorestown, NJ). We imaged the feather cross-sections using a Gemini 500 field-emission scanning electron microscope (Carl Zeiss, Oberkochen, Germany). We acquired images using an in-lens SE detector at an accelerating voltage of 3–5 kV, and additionally imaged FM-stained feathers using an annular energy-selective BSE detector at an accelerating voltage of 5 kV.

### 2h. Micro-computed tomography

We collected three-dimensional micro-computed tomography (μCT) scans of FM-stained feathers on a Zeiss 520 Versa X-ray microscope (Carl Zeiss, Oberkochen, Germany) at the Cornell Institute of Biotechnology imaging facility. To immobilize the delicate barbs and barbules during imaging, we mounted washed and FM-stained feathers between two layers of transparent tape. We acquired volumetric datasets of barbs and attached barbules with 0.12 μm voxel size using a 40x objective at 100kV/9W, and reconstructed from 1601 projections (15 s exposure each) using the standard Zeiss reconstruction software. We used Dragonfly [90] to visualize 2D projections and the 3D volume.

### 2i. Melanosome morphology analysis

We analyzed SEM images in Fiji v2.14.0 [91]. We could readily identify individual melanosomes as round-to-oblong bodies roughly 0.5–1 μm in size, and quantified their morphology by manually fitting the two-dimensional projection with an elliptical selection tool. We approximated the morphology and size of each melanosome using the aspect ratio and area of the ellipse fit:

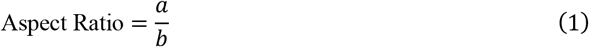

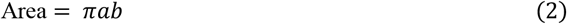

where *a* and *b* refer to the semi-major and semi-minor axis lengths, respectively. We used this method to likewise measure the morphology of individual melanin granules imaged in FM/BSE-SEM. Details on the statistical analysis used in our analysis of morphological data can be found in the supplemental information.

## 3. Results

### 3a. Fontana-Masson staining enables selective imaging of melanin deposits in the feather interior when coupled with backscatter scanning electron microscopy (FM/BSE-SEM) and micro-computed tomography (FM/μCT)

To selectively visualize melanin-rich structures (*i*.*e*., melanosomes) at different length scales against the keratinous matrix in the interior of feather barbs using SEM and μCT, we stained feathers with Fontana-Masson (FM) stain prior to imaging. The precipitation of elemental silver and gold on melanin, used in histopathology to darken melanosomes for optical microscopy, likewise allows melanin to be identified by the electron and X-ray contrast of these heavy elements relative to the organic matrix. (Fig. 2, 3). The difference in contrast between FM-stained white (*S. pileata* throat) and black feathers (*S. bouvreuil* crown) in FM/μCT (Fig. 2*a–b*) was striking: the white feather was completely devoid of melanin and the unstained keratin was barely visible compared to the bright melanin-rich regions in the black feather.

**Figure 2.**
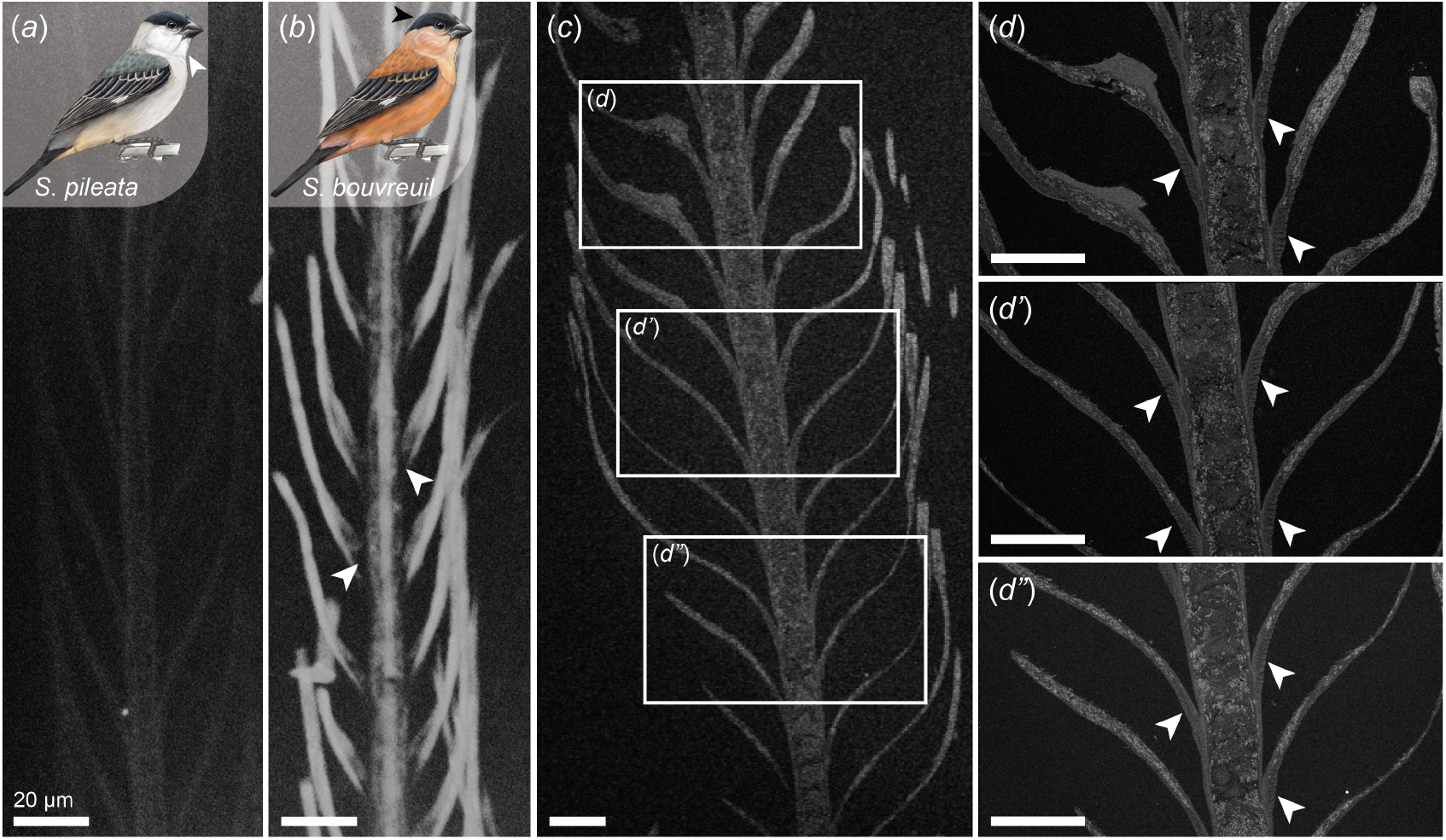
Melanin distribution visualized by FM/μCT and FM/BSE-SEM. (*a, b)* μCT maximum intensity projections of FM-stained white (*S. pileata* throat) and black (*S. bouvreuil* crown) feathers. Melanin in the black feather is highlighted by concentrated silver deposits clearly visible against the dark matrix. The single bright spot in the white feather is likely a nonspecific staining artifact. (*c–d”*) FM/BSE-SEM highlights individual melanosomes in the barbs and barbules of a black *S. bouvreuil* crown feather, corroborating the distribution visible in FM/μCT with melanin visible throughout the barb matrix, and dense deposits filling all but the proximal ~10 μm of each barbule (white arrows). Higher magnification views of (*d”*) can be seen in Figure 3(*b’–c’*). All scale bars 20 μm.

**Figure 3.**
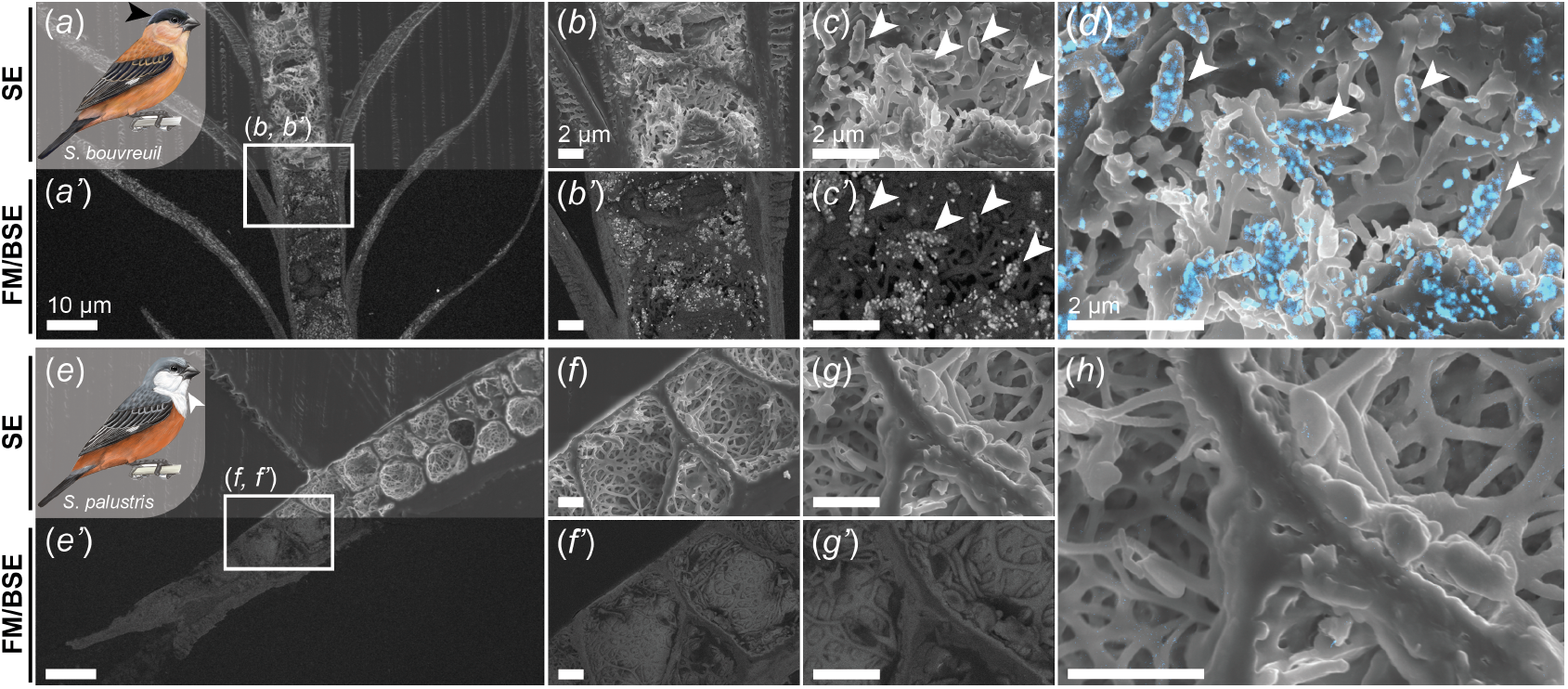
Melanin visualized by FM staining in SEM. (*a, a’*) Longitudinal section of a black *S. bouvreuil* crown feather, imaged in SE-SEM (top half) and FM/BSE-SEM (bottom half). (*b, b’*) SE-SEM and FM/BSE-SEM views of the region indicated by the white rectangle. (*c, c’*) Higher magnification images of the same region. (*d*) Brightness-thresholded false color overlay of the FM/BSE-SEM image in (*c*) onto the corresponding SE-SEM view in (*c’*). Arrows indicate melanosomes visible in both views. (*e, e’*) Similar longitudinal section of a white *S. palustris* throat feather with corresponding views (*f*–*g’*) showing an absence of any melanosomes (SE-SEM) or melanin (FM/BSE-SEM). (*h*) Overlay of the images in (*g*) and (*g’*) using the same threshold as in (*d*).

FM/μCT and FM/BSE-SEM (Fig. 2*b* and 2*c*–*d*, respectively) both revealed that melanin was distributed throughout the barbs and barbules of the pigmented feather. Imaging feather interiors with correlated SE-SEM and FM/BSE-SEM views (Fig. 2–3, S1) showed the presence of melanin even when melanosomes could not be discerned amid the convoluted keratin matrix. Individual melanosomes could be identified in the barb in SEM, and the barbules appeared densely packed with melanin except for the proximal 10 μm of each barbule, which was devoid of pigment (Fig. 2, white arrows). The SEM cross-section (Fig. 2*d–d”*) revealed that the melanin-free region, which anchors each barbule to the barb, consists of dense keratin contiguous with the outer cortex of the barb.

We imaged individual FM-stained melanin deposits within the barb with spatially correlated secondary electron (SE-SEM) and backscattered electron (FM/BSE-SEM) views. A comparison of longitudinal sections of representative black (*S. bouvreuil* crown; Fig. 3*a–c’*) and white (*S. palustris* throat; Figure 3*e–g’*) feathers confirmed that the elongated, 0.5–1 μm long structures identified in SE-SEM images contained melanin and were indeed melanosomes, further corroborated by their absence in the unpigmented feather. We observed that the silver stain did not uniformly fill each melanosome, but rather appeared in discrete, granular deposits. These granules were much smaller than a single melanosome, and approximately ellipsoidal with median major and minor axis lengths of 115.54 nm (interquartile range (IQR): 95.55–142.17 nm) and 81.51 nm (IQR: 69.65–95.97 nm), respectively, with a corresponding median aspect ratio of 1.372 (IQR: 1.217–1.607). While granules seen in FM/BSE-SEM generally coincided with melanosomes in the corresponding SE-SEM image, some granules appeared scattered in the keratin matrix; it is unclear whether these granules were truly free-standing, or were displaced by damage to melanosomes during sectioning.

### 3b. Melanosomes from crown feathers differ morphologically from those in other body plumage patches

We quantified melanosome morphology in Capuchino feathers by a manual elliptical fit of individual melanosomes in SE-SEM images (Fig. 4, S2–S3). Comparing melanosome aspect ratio and area, we found that crown feather melanosomes were largely distinct from those found in throat, dorsum/rump, and belly patches. Interestingly, while males of all the Capuchino species studied had a variety of patch coloration patterns, their crown feathers were invariably either black or gray (note: our collection included crown feathers for only eight of the twelve species, see Fig. 4 and Table S1). Crown feather melanosomes were elongated in all species except *S. iberaensis*, in which they were round. The elongated melanosomes were not homogeneous but rather covered a range of morphologies, from *S. pileata* with a median aspect ratio of 2.39 (IQR: 2.07–3.13) to *S. minuta* crown melanosomes with a median of 3.94 (IQR: 3.50–4.28). *S. iberaensis* crown melanosomes were nearly spherical, with a median aspect ratio of 1.20 (IQR: 1.16–1.30). We found that having oblong melanosomes was a distinguishing feature of the crown patch itself and not linked to black or gray feather color: feathers from the belly, throat, and dorsum/rump patches had round melanosomes regardless of color (with the exception of *S. nigrorufa* rump, see below). Melanosomes in crown feathers were also significantly larger (median: 0.39 μm^2^, IQR: 0.31–0.48 μm^2^) than those from the belly (median: 0.12 μm^2^, IQR: 0.09–0.20 μm^2^; *U* = 4726, *r* = 0.54, *p* < 0.00001), throat (median: 0.20 μm^2^, IQR: 0.15–0.26 μm^2^; *U* = 3425, *r* = 0.585, *p* < 0.00001), and dorsum/rump patches (median: 0.23 μm^2^, IQR: 0.11–0.39 μm^2^; *U* = 3711, *r* = 0.37, *p* < 0.00001). Interestingly, the larger area of crown melanosomes was not simply due to elongation in one direction: crown melanosomes had simultaneously longer long axis (*a*) lengths and shorter short axis (*b*) lengths (Fig. S3) than melanosomes from all other patches (*a* comparison: *U* = 2808, *r* = 0.46, *p* < 0.00001; *b* comparison: *U* = 15988, *r* = 0.13, *p* = 0.0003).

**Figure 4.**
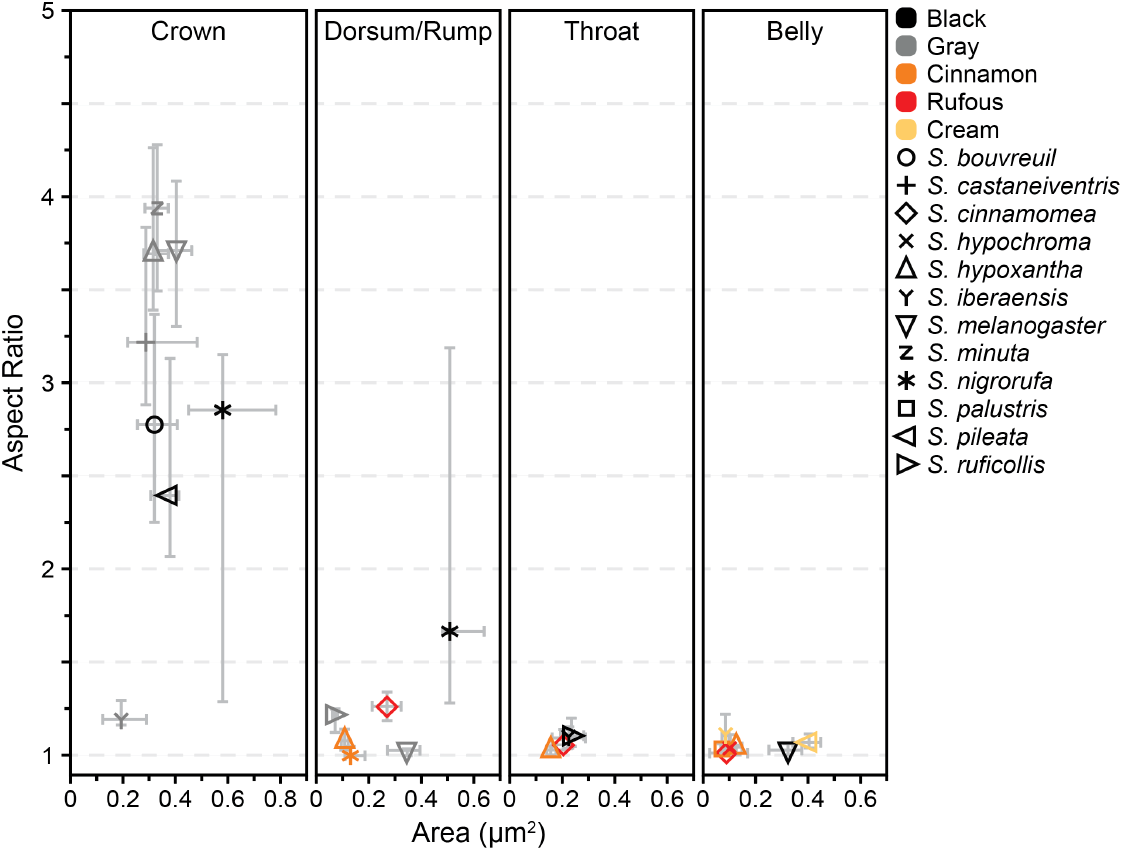
Quantification of Capuchino melanosome morphology. Each data point represents the median aspect ratio and area of all melanosomes measured in the indicated species and patch; error bars indicate the interquartile range (IQR).

The crown was the only patch that was exclusively EM-dominant (*i*.*e*., black or gray) across all Capuchino species. Therefore, we used crown feathers to compare patch-specific differences between black and gray feathers, and non-crown patches to compare the melanosome morphologies of PM-dominant (cinnamon/rufous) feathers to the EM-dominant ones. In the ventral patches (throat and belly), where black and gray feathers had nearly spherical melanosomes, we found that EM-dominant melanosomes were typically larger than PM-dominant melanosomes (Fig. 4; *U* = 21514.5, *r* = 0.54, *p* < 0.00001). The difference between black and cinnamon/rufous feathers was most striking in the belly patch, where melanosomes from black feathers had a median area of 0.33 μm^2^ (IQR: 0.25–0.38 μm^2^), much larger than those from cinnamon (0.12 μm^2^, IQR: 0.09–0.15 μm^2^; *U* = 964.5, *r* = 0.68, *p* < 0.00001) and rufous (0.11 μm^2^, IQR: 0.07–0.13 μm^2^; *U* = 155, *r* = 0.78, *p* < 0.00001) feathers. On the other hand, in the rump patch (the only dorsal patch in which cinnamon or rufous feathers were present), we found the area distributions of melanosomes from gray and rufous feathers to largely overlap. Cream feathers had more variability in melanosome size: while *S. pileata* belly melanosomes had the largest melanosomes (median area: 0.41 μm^2^, IQR: 0.34–0.45 μm^2^) of any ventral-aspect feathers, *S iberaensis* belly melanosomes were conversely among the smallest (median area: 0.09 μm^2^, IQR: 0.07–0.10 μm^2^).

The dorsum and rump patches of most species were the same color, and were for this reason grouped together and indicated as dorsum/rump. Only *S. nigrorufa* presented an exception by having distinct colors (black dorsum, rufous rump); the feathers from these two patches were therefore categorized separately. *S. nigrorufa* dorsal feather melanosomes (black; Fig. 4) were morphologically closer to those found in crown feathers, with a mix of oblong and round melanosomes with a median AR of 1.67 (IQR: 1.28–3.19) and area of 0.51 μm^2^ (IQR: 0.48–0.64 μm^2^), while its rump feathers (rufous) had small, round melanosomes with a median AR of 1.09 (IQR: 1.05–1.16) and area of 0.10 μm^2^ (IQR: 0.08–0.15 μm^2^).

### 3c. Feathers from dorsal patches have higher melanin content than ventral patches

Melanin composition [23] was largely predictive of feather color between black/gray and cinnamon/rufous feathers (Fig. 5, Table S2). The majority of the melanin content of cinnamon and rufous feathers consisted of PM, ranging from 70% PM (*i*.*e*., 30% EM) in rufous *S. cinnamomea* dorsum/rump feathers to 87% PM in cinnamon *S. cinnamomea* throat feathers. Cinnamon and rufous feathers had similar absolute PM content, ranging from 47.0 to 122.5 μg/mg and 46.7 to 107.9 μg/mg, respectively. Within the throat, belly, and dorsum/rump patches (the only patches that could have either cinnamon or rufous feathers), however, we found that cinnamon feathers had a higher BZ-PM:BT-PM pheomelanin subunit ratio (Fig. S4). The melanin content of black and gray feathers consisted predominantly of EM, ranging from 52% EM (black *S. ruficollis* throat) to 93% EM (black *S. nigrorufa* crown). The percent EM content of cream feathers was within the range of EM-dominant coloration, however, they had much less total melanin (18–20 μg/mg), particularly compared to the melanin content of all other belly feathers which ranged from 65.7 μg/mg to 92.7 μg/mg.

**Figure 5.**
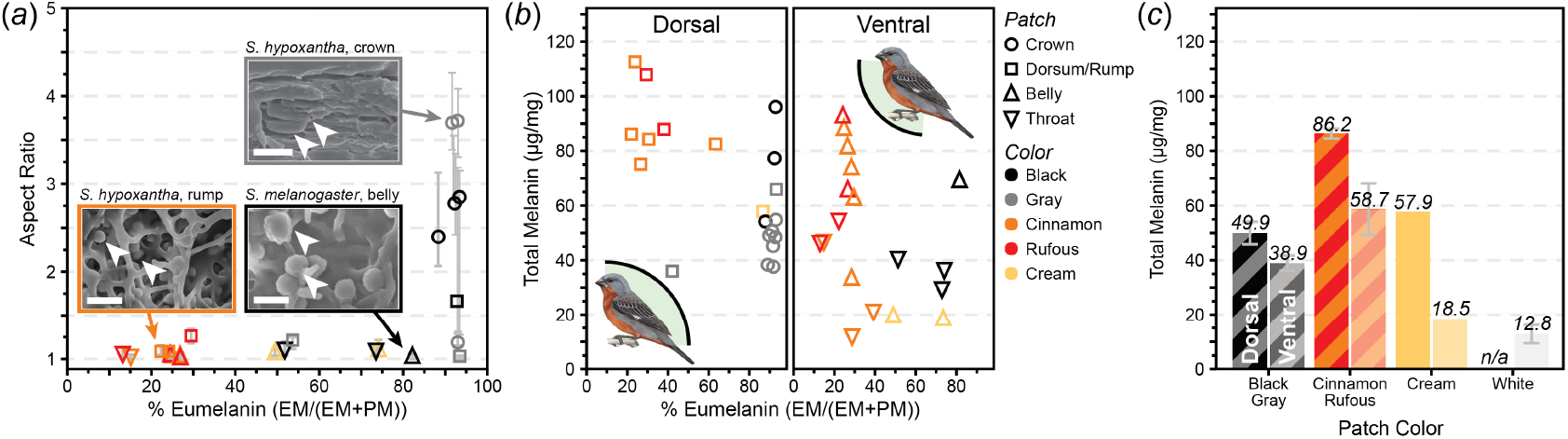
(*a*) Median melanosome aspect ratio plotted against EM content as a percentage of total melanin, grouped by species (not indicated) and patch (marker shape) with error bars indicating IQR. Insets show the range of melanosome morphologies seen across species and patches; all error bars 1 μm. (*b*) Pigment data from Estalles *et al*. (2022) shown as a plot of the total melanin and percentage EM content, grouped by anatomic aspect [23]. (*c*) Summary of pigment data further grouped by dominant melanin type to compare the melanin content of dorsal and ventral feathers in each coloration type.

Crown feathers differed from other patches in their EM content. As they were exclusively gray or black, all crown feathers were almost entirely composed of EM, with a minimum of 88% EM in black *S. pileata* crown (Fig. 5*a*). The variability in EM content was accordingly lower in crowns than all other patches: all crown feathers fell within five percentage points of each other, while the EM content of black and gray non-crown patches spanned a range of 30 percentage points across species.

In addition to these patch-specific differences, we also grouped the patches by anatomic aspect, *i*.*e*., dorsal (crown, dorsum, and rump) or ventral (belly and throat), and compared the EM content between the two groups. This grouping can be useful in identifying broader patterns between patches that are anatomically distinct, yet still share functional similarities: for example, by grouping the patches that receive higher sun exposure (dorsal) and those that are more visible during mating displays (ventral). Comparing these groups (Figs. 5*b*–*c*), we found that cinnamon and rufous dorsal feathers had significantly more total melanin than cinnamon and rufous ventral feathers (p = 0.0047). Black and gray dorsal feathers also generally had more total melanin than corresponding ventral feathers, but the difference was not statistically significant (p = 0.3258), and although cream patches were not numerous enough to be compared statistically, they likewise followed the same pattern.

## 4. Discussion

To investigate the relationship between pigment content, melanosome morphology, and feather coloration in the Capuchino Seedeaters, we compared melanin composition (from HPLC, reported in Estalles *et al*. 2022) and melanosome dimensions (measured from SE-SEM) across species and plumage patches. Capuchino feather colors were categorized into four broad color groups with distinct pigmentary and/or morphological parameters:

i. White feathers lacked melanin or melanosomes; this coloration instead typically arises from diffuse light scattering by the keratin matrix, but without the interference inherent to structural color. The neutral white color of unpigmented feathers suggests that the matrix likely did not affect feather coloration.
ii. Black and gray feathers were characterized by high EM content and larger melanosomes (relative to cinnamon and rufous feathers). Melanosome morphologies were variable but patch-specific, ranging from highly elongated in the crowns of most species, to nearly always round outside the crown patch. Black feathers could not be distinguished from gray ones solely by their melanosome morphology or pigment content: both colors could arise from overlapping ranges of melanosome morphologies and melanin content.
iii. Cinnamon and rufous feathers accordingly had high PM content and smaller, much less morphologically variable melanosomes. As with black and gray feathers, we did not observe clear morphological differences between the consistently round melanosomes of cinnamon and rufous feathers. We nevertheless found that when controlling for patch location, cinnamon feathers had a higher BZ-PM to BT-PM ratio than rufous feathers, suggesting that difference between cinnamon and rufous colors may arise in part from different pigment chemistries. This trend, however, did not hold for comparisons across different patches, so it remains unclear to what extent the PM subunit ratio contributes to the differentiation of cinnamon and rufous Capuchino feather colors.
iv. Cream feathers were only present in two species in our collection, and had round melanosomes with considerable interspecific variability in size: *S. iberaensis* had among the smallest and *S. pileata* had the largest of all the round melanosomes we measured. Cream feathers had a similar EM content to black feathers, but very little total melanin relative to all other pigmented feathers; taken together, the pigment content and the melanosome morphology suggest that cream feathers constitute a unique color in Capuchinos, and not simply a melanin-deficient variation.

Although species- and patch-specific patterns emerged in our analysis, we did not find any trends in melanosome morphology or pigment content that could reliably differentiate feather colors in a general manner independent of patch or species. Analysis of sectioned feathers showed that EM-dominant (black or gray) feathers in crown patches predominantly contained oblong melanosomes that were generally larger than the round melanosomes found in the black or gray feathers of the remaining patches.^1^ While melanosome elongation has been associated with higher EM content through the effects of a PMEL17 scaffold [35] we found several exceptions, notably *S. iberaensis* crown and *S. melanogaster* dorsum/rump feathers, with a very high proportion of EM yet exclusively spherical melanosomes. Therefore, as morphology did not account for the color differences in gray and black crown feathers, it remains unknown why the crown melanosomes of most Capuchino species shared an oblong morphology, and how comparable colors could also be achieved through round melanosomes (Fig. 5, S5).

The large variability in melanosome morphologies (even when grouped by species and patch), and the lack of straightforward correlations to plumage color, are nevertheless interesting features of the Capuchino model system. The observation of black feathers with round melanosomes in particular contradicts the generally accepted association of dark EM-dominant coloration with elongated melanosomes, and reddish PM-dominant coloration with round melanosomes [40,92,93]. In the Li *et al*. (2012) dataset [40] of melanosome morphologies across 127 extant bird species, all 45 species with black feathers have elongated melanosomes with mean aspect ratios above 2.0 (Fig. S6); on the other hand, four of the twelve Capuchino species had black feathers with melanosome aspect ratios between 1.0 (spherical) and 2.0. Quadratic discriminant analysis of the published dataset has shown that dark EM feather colors could still be differentiated even when melanosome dimensions showed no clear correlation, by considering distribution parameters like the coefficient of variation (CV) and skewness. The poor agreement of black and gray Capuchino melanosomes with this model (Fig. S6) was due in part to the correlation of aspect ratios with patch location rather than feather color as would be expected, nevertheless this shows that bird feather coloration cannot be fully explained by melanosome morphology and complicates efforts to generalize such a relationship to different species and particularly pigment types.

The association of melanosome morphology and pigment content with plumage patches virtually irrespective of feather color was also notable and may offer insights, albeit tentative, into the factors affecting the divergence of the different patch coloration patterns observed in the Capuchinos. The lack of color variation in crown feathers (particularly relative to their highly variable melanosome morphologies between and within species), along with the higher pigment density and EM content seen in dorsal patches, may suggest that selective pressures favored dark, EM-rich coloration. The Capuchinos are migratory and inhabit sunny grassland habitats [30], and the features shared by dorsal feathers are consistent with an adaptation to protect upward-facing feathers (and perhaps tissues) from UV damage [1,70].

Conversely, the presence of brightly colored (and highly variable across species), PM-dominant or melanin-deficient feathers in male Capuchino belly and throat patches supports the hypothesis that ventral feathers play an important signaling role in mate recognition and sexual selection [30,32]. Producing feather pigments, particularly in seasonally molting species like the Capuchinos, can be resource-intensive [1,94], and the coexistence of bright PM-rich and dark EM-rich feathers in different plumage patches of the male mating coloration points to a balance between the selective pressures of mating and photoprotection. The localization of pigments with opposing properties in dorsal and ventral patches suggests careful resource allocation to compromise between the simultaneous need for bright, mate-signaling and dark, UV-protective feathers. Nevertheless, evidence for this dual role remains circumstantial: a reflectance study of *S. hypoxantha* and *S. iberaensis* [32] did not find consistent differences in UV reflectance between dorsal and ventral patches in either species, and confidently assigning functions to Capuchino plumage patches and colorations would require more rigorous analysis.

We describe a novel application for Fontana-Masson (FM) stain to visualize melanin distribution and individual melanosomes at high resolution. Due to the ability of melanin to reduce silver ions, which precipitate as dark brown deposits of elemental silver, FM staining has traditionally been applied in histology (*i*.*e*., optical microscopy) to diagnose melanoma [95,96] and identify melanin-producing microorganisms like *Cryptococcus neoformans* in fungal infections [97–101]. FM staining has recently been used with μCT [102] to map melanin distribution in zebrafish, however, most commercially available μCT systems do not have sufficient resolution to distinguish individual melanosomes. To our knowledge, FM staining has never been used to image individual melanin granules at the higher resolution (sub-nanometer) enabled by scanning electron microscopy. Here, we show that FM staining can provide selective contrast enhancement in a high-resolution application in BSE-SEM, and label individual melanin granules (Fig. 3) with high specificity. This accessible, off-the-shelf solution allows melanosomes to be unambiguously identified in SEM, benefiting not only from the ability to image with secondary-electron (topography) and backscatter (Z-contrast) detectors, but also from more straightforward sample preparation and higher throughput than TEM.

As a proof-of-principle, we emphasize the flexibility of this staining technique: FM stain is readily commercially available and can be easily applied to whole feathers, and its high specificity for melanin (and by extension melanosomes) was evident in both FM/μCT and FM/BSE-SEM. The high contrast of melanin-bound silver against the unstained organic matrix, combined with the relatively large escape depth of backscattered electrons, make FM/BSE-SEM a valuable complement to conventional SE-SEM by allowing melanosomes to be resolved even in challenging settings where they may be partially obscured by other matrix structures (Fig. 3). Although FM/μCT and FM/BSE-SEM were done on separate feathers in this work, we propose that the two techniques could be used sequentially on the same feather as a means of correlative imaging. Combining FM/BSE and FM/μCT in this way enables multiscale characterization of melanin distribution, from individual melanosomes to entire feather barbs.

Here, FM/BSE and FM/μCT were used to visualize the distribution of melanin in Capuchino feathers and revealed densely packed melanosomes in barbules as well as many scattered throughout the keratinous matrix. As melanosomes have been linked to mechanical stiffening of feathers [1,103], the melanin-free proximal region that connects each the barbule to the cortex may play a role in ensuring flexibility by functioning as a soft joint compliant enough to withstand stresses such as the disentangling of barbules during preening.

## 5. Conclusion

In this study, we characterized color production across the Capuchinos through a combined approach examining differences in both pigment production and melanosome morphology. We identified patch-specific patterns in melanosome morphology and melanin pigment content: melanosomes from black and gray EM-dominant feathers had highly variable morphologies across different body patches, with the crown feathers predominantly containing elongated melanosomes while the belly, throat, and dorsum/rump patches generally had round melanosomes irrespective of plumage color. We found that dorsal feathers, which were either gray or black in most Capuchino species, consistently had higher percent EM content and a greater total melanin density than ventral feathers of the same color. Patterns in the variability of melanosome morphology and patch coloration may suggest a possible interplay of evolutionary pressures in the Capuchino speciation, but further work is needed to test the strength of these inferences. Nevertheless, the morphological, pigmentary, and coloration patterns we describe suggest that melanosomes play a central role in the emergence of different plumage colors in the Capuchino Seedeaters.

Finally, our work describes the use of Fontana-Masson stain to selectively visualize melanin and melanosomes within feathers at different length scales with FM/μCT and FM/BSE-SEM. We hope this application of FM to high-resolution imaging will be useful for the community of researchers interested in melanin, melanosomes, and their consequences for feather coloration.

Our combined approach of pigmentary and morphological characterization reveals how variations in both melanin content and melanosome morphology relate to the complexity of plumage coloration patterns. Our findings emphasize the importance of comprehensive, multi-scale analysis in interpreting the functional and evolutionary roles of melanin-based color.

## Data Accessibility

All data used in this work is available at Dryad (DOI: 10.5061/dryad.8kprr4z1v).

## Declaration of AI use

We have not used AI-assisted technologies in creating this article.

## Authors’ contributions

S.M.S.: Conceptualization, formal analysis, investigation, methodology, visualization, writing – original draft, and writing – review & editing. O.G.R.: Conceptualization, formal analysis, funding acquisition, investigation, methodology, and visualization. D.L.B.: Investigation, methodology, formal analysis, and writing – review & editing. C.B.D.: Conceptualization, methodology, and writing – review & editing. T.J.P.: Investigation, methodology, and writing – review & editing. L.F.S.: Resources and writing – review & editing. S.P.T.: Resources and writing – review & editing. L.A.E.: Conceptualization, funding acquisition, project administration, supervision, and writing – review & editing. L.C.: Conceptualization, funding acquisition, project administration, resources, supervision, writing – original draft, and writing – review & editing.

## Conflict of interest declaration

The authors declare no competing interests.

## Funding

This work was supported through the Cornell Lab of Ornithology Athena Fund (to O.R.), the College of Engineering Learning Initiatives undergraduate research award (to D.L.B.), and award DEB-2232929 from the National Science Foundation (to L.C. and L.A.E.). C.B.D. was supported by NSF ADBC TCN grant number 1700908 (oVERT). Museum collections were supported by the Fundação de Amparo à Pesquisa do Estado de São Paulo FAPESP (São Paulo Research Foundation), grants 2017/23548-2 and 2022/11650-6 (to L.F.S.). SEM was done at the Cornell Center for Materials Research shared instrumentation facility, and μCT data was acquired through the Cornell Institute of Biotechnology Imaging Facility, with NIH 1S10OD012287 funding for the Zeiss-Xradia Versa 520 X-ray microscope.

## Acknowledgements

All Capuchino illustrations by Jillian Ditner. We thank the Ornithology collections of the Centro de Ecología Aplicada del Litoral (CECOAL; Corrientes, Argentina) and the Museu de Zoologia da Universidade de São Paulo (São Paulo, Brazil) for the feathers used in this study. We also thank John Grazul (Cornell Center for Materials Research) for his help in designing the embedding protocol and his expertise in SEM sample preparation, and the staff of the Cornell University College of Veterinary Medicine Animal Health Diagnostic Center for FM-staining of whole feathers. Finally, we would like to thank Jennie A.M.R. Kunitake and Vanya Rohwer for their guidance and many valuable discussions of experimental design and data analysis.

## Footnotes

1 Because we calculated melanosome dimensions from SEM images, it is important to note that the projection of 3D objects in a 2D image may introduce certain measurement bias: scattered oblong melanosomes, for example, could by foreshortening give a false appearance of different morphologies depending on their individual orientation relative to the focal plane. Several Capuchino crown feathers had very broad melanosome aspect ratio distributions ranging from 1 to ~3–4, and we anticipate that some portion of this variation may arise from such imaging bias. The observation of one species (*S. iberaensis*) with exclusively round crown melanosomes and several in which no round-appearing melanosomes were seen, along with the general agreement between the melanosome dimensions reported here and the those measured in other birds by Li *et al*. [43,44], nevertheless suggests that the observed variability in crown melanosome aspect ratios was largely reflective of their true morphology.

## Notes

### Competing Interest Statement

The authors have declared no competing interest.

