## Supplementary Material for "A multiscale physicochemical characterization of melanin-based feather coloration in the rapid Capuchino Seedeater radiation"

**Electronic Supplementary Material for "A multiscale physicochemical characterization of melanin-based feather coloration in the rapid Capuchino Seedeater radiation"**

Stephan M. Sutter*, Olivia G. Rogers, Daniel L. Bilezikian, Casey B. Dillman, Teresa J. Porri, Luís Fábio Silveira, Sheela P. Turbek, Lara A. Estroff*,**, Leonardo Campagna*,**

**These authors contributed equally to the paper

**Fontana-Masson Staining Protocol** (Cornell University College of Veterinary Medicine Animal Health Diagnostic Center)

Solutions:

*(Stock) Ammoniacal Silver Nitrate:*

a. Add 10.0 g silver nitrate ($AgNO_3$) to 100.0 mL deionized (DI) water, yielding a 10% $AgNO_3$ solution.
b. To 95 mL of this solution, add concentrated (28%) ammonium hydroxide ($NH_4OH$) dropwise until solution is clear with no precipitate.
c. Add the remaining 10% $AgNO_3$ solution dropwise and refrigerate overnight.

*(Working) Ammoniacal Silver Nitrate:*

Dilute 10.0 mL of stock solution in 30.0 mL of DI water.

*0.2% Gold Chloride:*

Dilute 10.0 mL of 1% aqueous gold chloride ($AuCl_3$) in 40.0 mL of DI water.

*5% Sodium Thiosulfate:*

Dissolve 25.0 g of sodium thiosulfate ($Na_2S_2O_3$) in 500.0 mL of DI water.

FM Staining Protocol:

1. Hydrate feathers with deionized (DI) water
2. Submerge in working ammoniacal silver nitrate solution at 58°C for 45 min. to 1 hour. Remove when feathers appear light brown.
3. Rinse with DI water
4. Submerge in 0.2% gold chloride solution for a few seconds until feathers turn from light brown to gray
5. Rinse with DI water
6. Submerge in 5% sodium thiosulfate solution for 1 min.
7. Rinse well with DI water for 1 min.

**Statistical Analysis of Morphological Data**

Because many of the parameter distributions measured were not normal, we report summary statistics as median and interquartile range (IQR) where appropriate. We accordingly made pairwise comparisons with a two-tailed Mann-Whitney $U$ test and a $P < 0.05$ significance threshold. For large samples where the $U$-statistic was approximately normally distributed, we also calculated the standardized $z$-score by dividing the difference between the $U$-statistic and the mean by the standard deviation. From the $z$-score, we further report the effect size $r$ as the $z$-transformed statistic divided by the square root of the total sample size; distribution differences are approximated using the ranges $0.1 \leq r < 0.29$ (small difference), $0.3 \leq r < 0.49$ (moderate difference), and $0.5 \leq r \leq 1$ (large difference).

**Table S1. Summary of *Sporophila* feather SEM imaging**

| Species | Patch | Color | Individuals | Feathers | Images | Melanosomes |
|---|---|---|---|---|---|---|
| *S. bouvreuil* | Crown | Black | 2 | 2 | 5 | 67 |
| *S. castaneiventris* | Crown | Gray | 3 | 3 | 4 | 39 |
| *S. cinnamomea* | Throat | Cinnamon | 1 | 1 | 4 | 37 |
| | Belly | Rufous | 1 | 1 | 4 | 24 |
| | Rump | Rufous | 1 | 1 | 3 | 37 |
| *S. hypochroma* | Belly | Rufous | 1 | 1 | 3 | 30 |
| *S. hypoxantha* | Crown | Gray | 1 | 1 | 4 | 41 |
| | Throat | Cinnamon | 2 | 2 | 5 | 37 |
| | Belly | Cinnamon | 3 | 3 | 6 | 126 |
| | Rump | Cinnamon | 1 | 1 | 3 | 34 |
| *S. iberaensis* | Crown | Gray | 5 | 5 | 9 | 34 |
| | Throat | Black | 7 | 9 | 22 | 161 |
| | Belly | Cream | 1 | 1 | 4 | 52 |
| *S. melanogaster* | Crown | Gray | 1 | 1 | 1 | 13 |
| | Belly | Black | 3 | 3 | 7 | 65 |
| | Rump | Gray | 2 | 2 | 5 | 47 |
| *S. minuta* | Crown | Gray | 1 | 1 | 2 | 55 |
| *S. nigrorufa* | Crown | Black | 1 | 1 | 5 | 20 |
| | Dorsum | Black | 1 | 1 | 3 | 15 |
| | Rump | Cinnamon | 1 | 1 | 2 | 24 |
| *S. palustris* | Belly | Cinnamon | 1 | 1 | 2 | 32 |
| *S. pileata* | Crown | Black | 1 | 1 | 2 | 26 |
| | Belly | Cream | 1 | 1 | 4 | 24 |
| *S. ruficollis* | Throat | Black | 1 | 1 | 6 | 20 |
| | Dorsum | Gray | 1 | 1 | 3 | 54 |

**Table S2. Melanin Content Quantification (in µg of pigment per mg of feather)**

| | Crown | | | | Rump | | | |
|---|---|---|---|---|---|---|---|---|
| **Species** | **Color** | **EM** | **BT-PM** | **BZ-PM** | **Color** | **EM** | **BT-PM** | **BZ-PM** |
| *S. bouvreuil* | Black | 71.59 | 5.64 | 0.13 | Cinnamon | 20.22 | 30.67 | 24.3 |
| *S. cinnamomea* | Gray | 41.5 | 3.77 | 0.06 | Rufous | 31.88 | 41.28 | 34.73 |
| *S. hypochroma* | Gray | 34.12 | 3.71 | 0.52 | Rufous | 33.74 | 29.82 | 24.41 |
| *S. hypoxantha* | Gray | 34.62 | 2.91 | 0.18 | Cinnamon | 27.13 | 32.06 | 53.38 |
| *S. iberaensis* | Gray | 45.14 | 3.19 | 0.19 | Cream | 50.2 | 6.94 | 0.71 |
| *S. melanogaster* | Gray | 51.03 | 3.71 | 0.15 | Gray | 61.52 | 4.05 | 0.4 |
| *S. nigrorufa* | Black | 89.53 | 6.46 | 0.12 | Cinnamon | 19.19 | 27.95 | 39.06 |
| *S. palustris* | Gray | 46.32 | 4.22 | 0.24 | Cinnamon | 26.07 | 20.64 | 37.66 |
| *S. pileata* | Black | 47.80 | 6.36 | 0.09 | Gray | 15.16 | 15.57 | 5.18 |
| *S. ruficollis* | Gray | 44.04 | 4.52 | 0.52 | Cinnamon | 52.48 | 12.07 | 18.02 |

| | Belly | | | | Throat | | | |
|---|---|---|---|---|---|---|---|---|
| **Species** | **Color** | **EM** | **BT-PM** | **BZ-PM** | **Color** | **EM** | **BT-PM** | **BZ-PM** |
| *S. bouvreuil* | Cinnamon | 9.58 | 10.64 | 12.98 | Cinnamon | 8.59 | 6.12 | 7.1 |
| *S. cinnamomea* | Rufous | 17.67 | 20.57 | 27.49 | Rufous | 6.16 | 16.9 | 23.63 |
| *S. hypochroma* | Rufous | 22.69 | 34.1 | 35.94 | Rufous | 12.39 | 15.13 | 27.53 |
| *S. hypoxantha* | Cinnamon | 21.81 | 19.45 | 40.22 | Cinnamon | 7.11 | 12.58 | 27.39 |
| *S. iberaensis* | Cream | 13.64 | 3.94 | 0.87 | Black | 21.74 | 7.07 | 0.82 |
| *S. melanogaster* | Black | 56.7 | 10.71 | 1.76 | Black | 27.51 | 8.3 | 1.27 |
| *S. nigrorufa* | Cinnamon | 18.73 | 18.19 | 25.39 | Cinnamon | 3.64 | 4.62 | 4.26 |
| *S. palustris* | Cinnamon | 22.04 | 22.3 | 43.8 | White | 4.67 | 1.1 | 0.19 |
| *S. pileata* | White | 9.65 | 7.07 | 2.84 | White | 7.71 | 4.15 | 0.95 |
| *S. ruficollis* | Cinnamon | 21.17 | 17.85 | 34.61 | Black | 21.09 | 13.91 | 5.82 |

*Note: Pigment content is represented by the respective amounts of eumelanin (EM) and pheomelanin (PM), the latter consisting of benzothiazine (BT-PM) and benzothiazole (BZ-PM) moieties. Data originally published in Estalles* et al. *(2022) [23].*

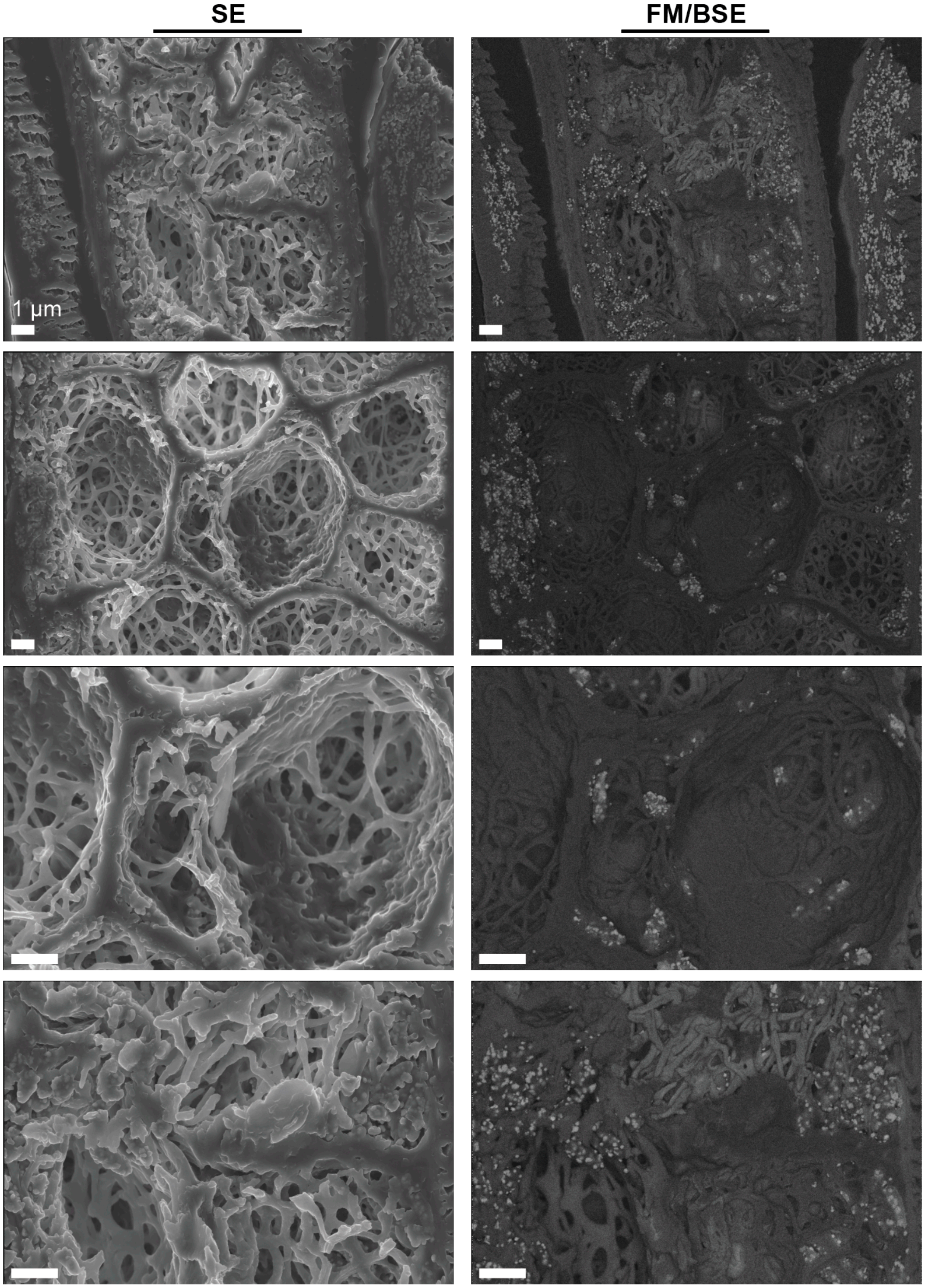


**Figure S1.** Spatially correlated SE-SEM (*left*) and FM/BSE-SEM (*right*) views of the interior of a black *S. bouvreuil* crown feather. Bright spots in FM/BSE-SEM identify melanin deposits even when individual melanosomes are not visible in SE-SEM. All scale bars 1 µm.

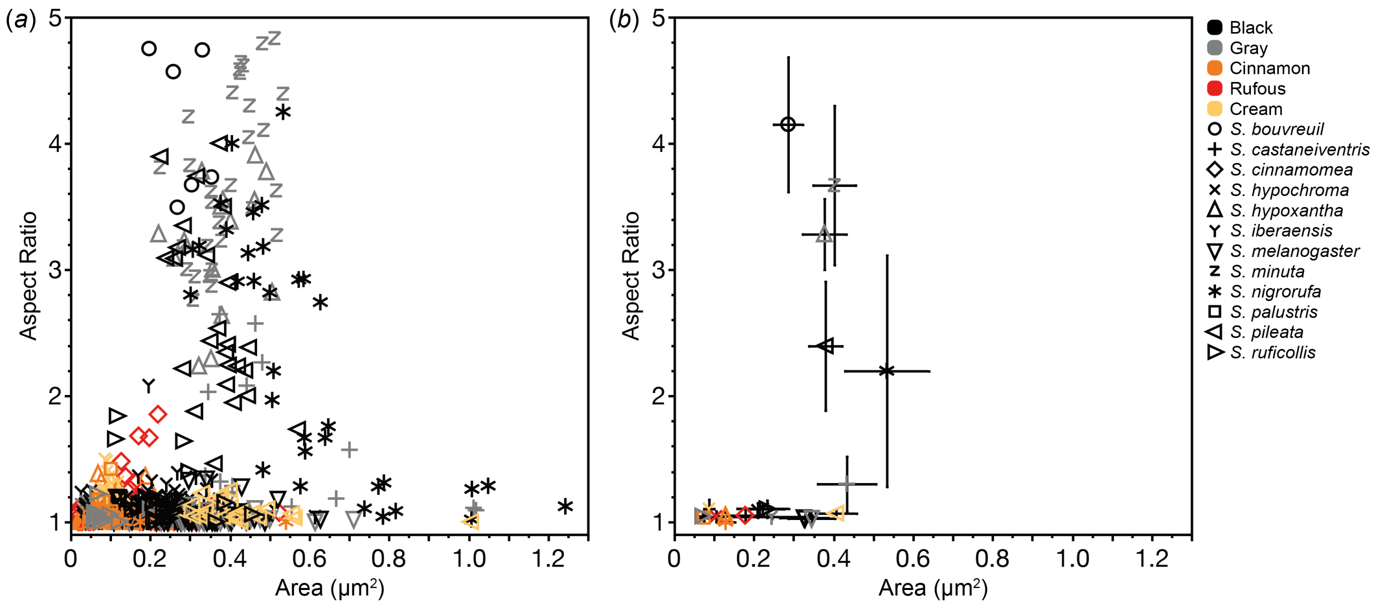


**Figure S2.** (*a*) Morphological metrics of all Capuchino species; each point corresponds to a single melanosome. (*b*) The same data grouped by species and color for clarity; points indicate the median with error bars showing median absolute deviation (MAD).

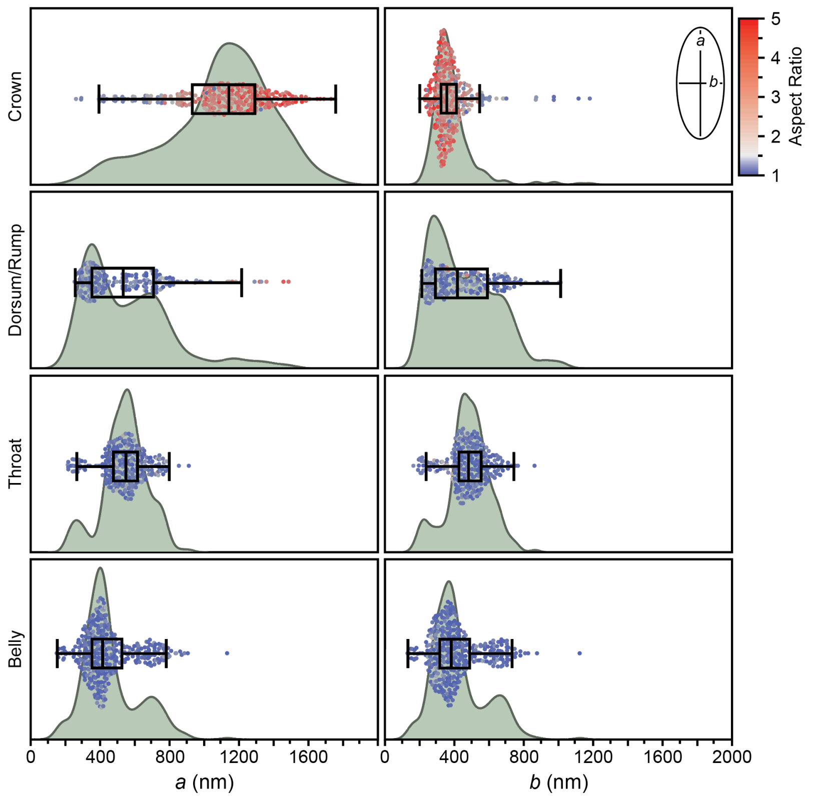


**Figure S3.** Distributions of *a* (major axis) and *b* (minor axis) lengths grouped by patch. Points are colored by aspect ratio, with the color scale centered at 1.5 to distinguish elongated and round melanosomes. Elongated melanosomes (red), primarily found in the crown, not only had long *a* lengths but also relatively short *b* lengths, especially compared to round crown melanosomes.

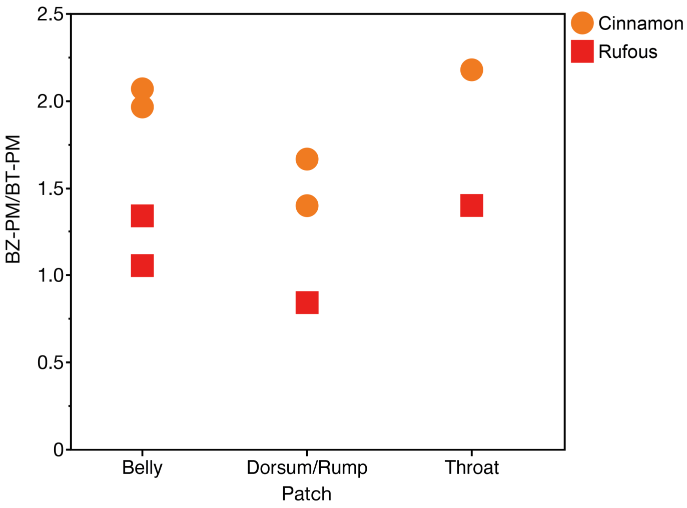


**Figure S4.** Comparison of PM composition, represented as the BZ:BT subunit ratio, between cinnamon and rufous feathers for species in which both colors appeared. In these cases, cinnamon feathers consistently had a higher BZ:BT ratio.

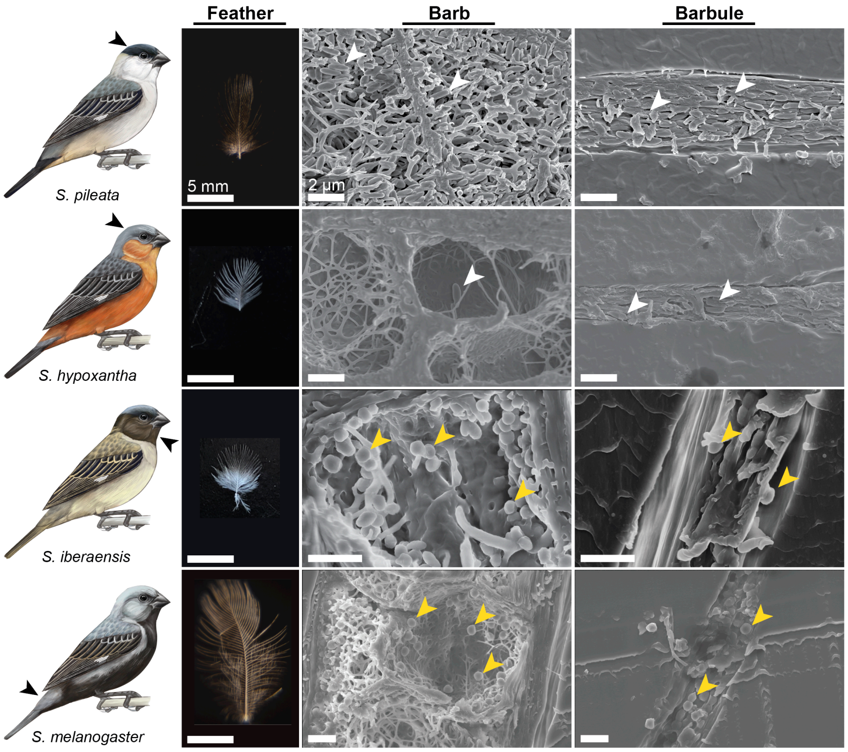


**Figure S5.** Optical microscopy (*left*) and representative barb (*center*) and barbule (*right*) cross-sections (SE-SEM) of a gray feather from each plumage patch, showing how different melanosome morphologies (*e.g.*, elongated crown melanosomes or round throat melanosomes) can result in the same feather color. White and yellow arrows indicate elongated and round melanosomes, respectively.

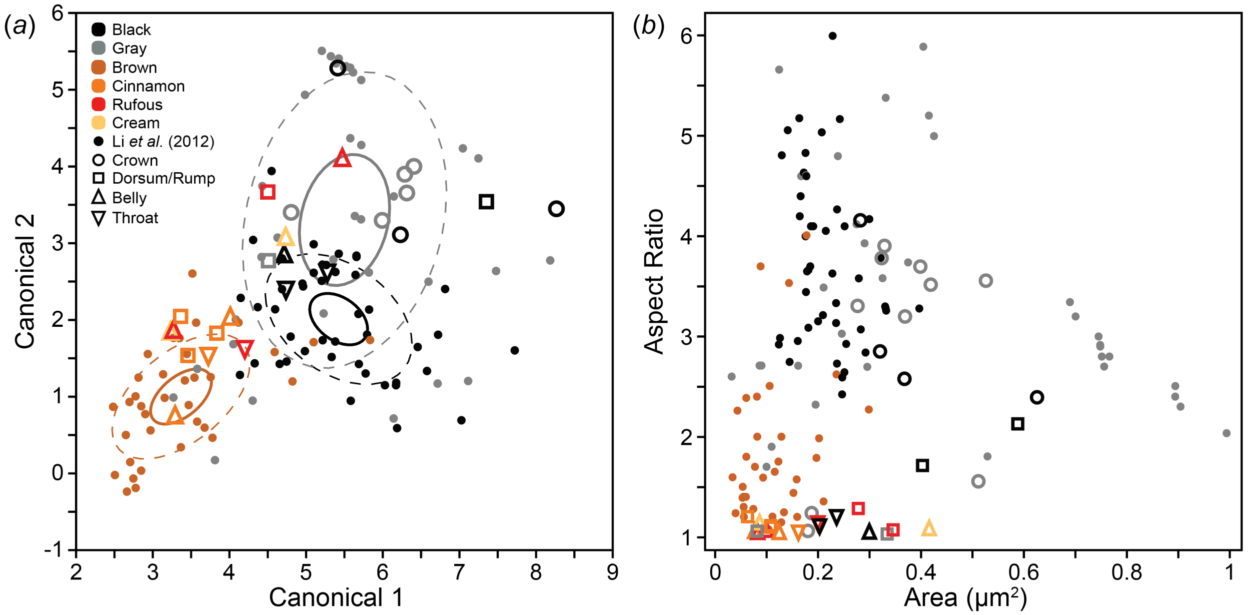


**Figure S6.** (*a*) Capuchino melanosomes (grouped by species and patch) overlaid onto the quadratic discriminant analysis (QDA) model from Li *et al.* (2012) [40]; only black, gray, and brown feathers from this dataset are shown for clarity. Solid ellipses indicate the 95% confidence bound for each color, while dashed ellipses indicate normal 50% contours. (*b*) Morphological parameters of the same data, with aspect ratio and area calculated from melanosome dimensions.
